# Do Memories of Inferred Visual Representations Guide Low-level Perception?

**DOI:** 10.1101/2025.03.07.641940

**Authors:** Ruoying L. Wang, William J. Harrison

**Affiliations:** School of Psychology, University of Queensland; School of Health, University of Sunshine Coast

**Keywords:** recognition memory, visual awareness, psychophysics, illusory perception

## Abstract

Prior knowledge shapes how we interpret and adapt to a dynamic environment. A crucial aspect of this process is the lifelong development of structured object representations, enabling meaningful and survival-relevant interactions with the external world. In this study, we investigated the extent to which memories of complex objects influence perception by refining visual details. To test this, we had participants hold in working memory Mooney images, a stimulus class that requires top-down processing to perceive hidden image structure. While holding a Mooney image in memory, participants performed a detection task, in which they had to detect an edge feature that appeared at selected locations of the illusory contours of the image. Participants completed this task twice, once before they were shown the hidden Mooney structure, and then after learning the hidden structure. In a signal detection framework, we assessed whether learning the hidden content altered participants’ sensitivity and response bias in detecting edge-feature targets. We found that, while holding object memories enhanced Mooney image disambiguation, it did not refine visual sensitivity. This dissociation — where categorical identification improves without corresponding perceptual refinement — suggests that memories of complex objects improve the overall understanding of ambiguous information independent of refining relevant visual details. These findings have important implications for the theory of recurrent processing, which has traditionally emphasised perceptual refinement in top-down feedback. Our results highlight how prior knowledge improves the perceived clarity of degraded visual information without necessarily improving precision in local feature detection.

---

Conceptual knowledge of visual objects is encoded in memory as semantically meaningful representations (Clarke & Tyler, 2015; DiCarlo et al., 2012; Xue, 2022). Imagine, for example, a baby learning what an apple is: each encounter with the fruit builds the baby’s understanding through the fruit’s appearance, feel, and taste. Such experiences accumulate over time so that, in the future, simply seeing the shape of an apple can trigger a system of complex understanding, giving rise to a variety of potential interactions, from eating to recognising it as part of healthy eating advice (Kragel et al., 2021; Squire & Wixted, 2011; Xue, 2022). But do these *abstract* associations merely guide cognition and behaviour, or might they also *refine* visual perception — perhaps altering the future processing of the same objects?

Research suggests that conceptual knowledge interacts with perception through working memory (Brady et al., 2008; Bruning & Lewis-Peacock, 2020; Csorba et al., 2022; Rademaker et al., 2019). Working memory, which is the capacity to temporarily hold information in mind, may influence perception because the two systems are thought to share neural resources (Bosch et al., 2014; Gayet et al., 2013, 2018; Rademaker et al., 2019; Teng & Kravitz, 2019; but see Harrison & Bays, 2018; and Xu, 2017). For example, holding a memory of a specific orientation can distort the perception of a subsequent stimulus (Teng & Kravitz, 2019). Similarly, the perception of masked geometric shapes is enhanced when related items are held in working memory (Gayet et al., 2013, 2016).

The apparent interaction between memory and perception extends beyond simple stimuli. Emerging evidence suggests that even complex object representations, traditionally considered the domain of long-term memory storage, can be actively maintained in working memory (Asp et al., 2021; Brady et al., 2008, 2016; Bruning & Lewis-Peacock, 2020; Hirschstein & Aly, 2023; Stokes et al., 2012). These findings suggest that stored object representations may not only passively exist in long-term memory but also actively shape ongoing perception (Clarke & Tyler, 2015; Hirschstein & Aly, 2023), raising the question of how this dynamic unfolds over time.

One possible underlying mechanism is **perceptual refinement** — the iterative improvement of visual representations through interactions between memory and perception (Bar, 2004; Bar et al., 2006). This process can be considered a corollary of the predictive coding framework, a theory proposing that the brain generates and updates its predictions about sensory inputs (Lee, 2002). In the context of object recognition, perceptual refinement is the idea that an initial coarse guess about an object’s identity is progressively refined by incorporating additional details (Bar, 2004; Bar et al., 2006; Hegde, 2008; Kveraga et al., 2007). For example, under conditions of uncertainty, expectations about an object are iteratively updated as more sensory information becomes available, leading to a more detailed and accurate representation (Boehler et al., 2008; Kirchberger et al., 2021; Polack & Contreras, 2012; Rajaei et al., 2019; Seijdel et al., 2021). This influential theory has shaped our understanding of object recognition and has inspired numerous computational vision models, where recurrent mechanisms are extensively implemented to enhance object recognition under occlusion (Pang et al., 2021; Rajaei et al., 2019; Wyatte et al., 2012; Kubilius et al., 2018; O’Reilly et al., 2013).

Despite the theoretical importance of recurrent processing, whether complex, meaningful, real-life objects are iteratively refined by perceptual systems remains underexplored (Asp et al., 2021; Brady et al., 2016). Much of the prior research has focused on simple memory items like lines, shapes, and orientations (Ester et al., 2009; Gayet et al., 2016; Rademaker et al., 2019). While some have shown that stimuli of real-world complexity influence perception, they have not explicitly measured whether the expectation of a complex object induces perceptual refinement. For example, Linde-Domingo et al. (2019) examined reaction times and neural activity during the retrieval of semantic labels (conceptual knowledge) and line drawings (perceptual details). Their findings suggest that the processing of semantic information precedes the retrieval of perceptual details, indicating that conceptual knowledge may guide visual processing. However, while reaction times were used, the study did not directly quantify changes in the quality of visual representations over time. Similarly, Asp et al. (2021) showed that memories of recognisable face images allowed a greater retrieval of image-target locations than their scrambled counterparts, suggesting an enhancement of working memory capacity. While their findings highlight the role of meaningful object representations in facilitating memory performance, they did not show *perceptual* refinement. To accurately quantify *changes in perception*, a rigorous psychophysics paradigm is required — a critical gap in the literature that our study aims to address.

## Investigating Perceptual Refinement Using Mooney Images

To isolate the influence of holding a complex object in memory on perception, an ideal behavioural paradigm should require continuous memory maintenance while participants engage in perceptual tasks. Studies of recurrent processing typically use backward masking, in which the presentation of a subsequent masking stimulus reduces the visibility of an initial target object. Backward masking may be effective for testing visual awareness (Gayet et al., 2018) because it forces the visual system to rely on partly visible or coarse features to make rapid categorical judgments. It does not, however, lend itself to quantifying the *perceptual implications* of retrieving object information in memory recurrently for at least two reasons. First, the reliance on a masking stimulus introduces low-level perceptual interference that can confound the effects of memory maintenance (Rademaker et al., 2019; Raymond et al., 1992; Robinson et al., 2019; Seiffert & Di Lollo, 1997). Second, masking paradigms intentionally restrict the time available for participants to fully retrieve complex, meaningful object memories, as the target is typically presented for only a brief interval (around 200 mms) before being disrupted. This is problematic because the disambiguation of conceptual knowledge is known to emerge on a longer timescale during retrieval (Asp et al., 2021; Clarke et al., 2011; Linde-Domingo et al., 2019; Teichmann et al., 2020). Conversely, delaying the masking stimulus risks allowing certain features to reach awareness, undermining the paradigm’s reliability for isolating memory-driven processes (Robinson et al., 2019). These limitations suggest that an alternative approach is needed to better elucidate the effects of memory on perception.

An alternative approach to backward masking is to use stimuli whose perceptual appearance strongly depends on an observer’s prior expectations. Teufel et al. (2018) developed a clever paradigm in which they exploited Mooney images to investigate the influence of object knowledge on low-level perception. Mooney images are (usually) modified photos that are reduced to coarse regions of black and white, lacking fine detail (Figure 1). To an observer, Mooney images first appear as images of random black-and-white blobs. However, the observer can learn the latent image structure, which fundamentally changes the image’s appearance when the visual system fills in missing information based on partial cues (Mooney, 1957). Teufel et al. (2018) found that when participants learned about the hidden content within these images, their ability to detect low-level visual details was enhanced relative to before they learned the hidden content. Detectability, therefore, depended on the observer’s prior knowledge of latent image structure. This result is consistent with the idea of recurrent processing, where higher-level expectations alter early sensory processing, thereby refining perceptual decisions (Bar et al., 2006).

**Figure 1.**
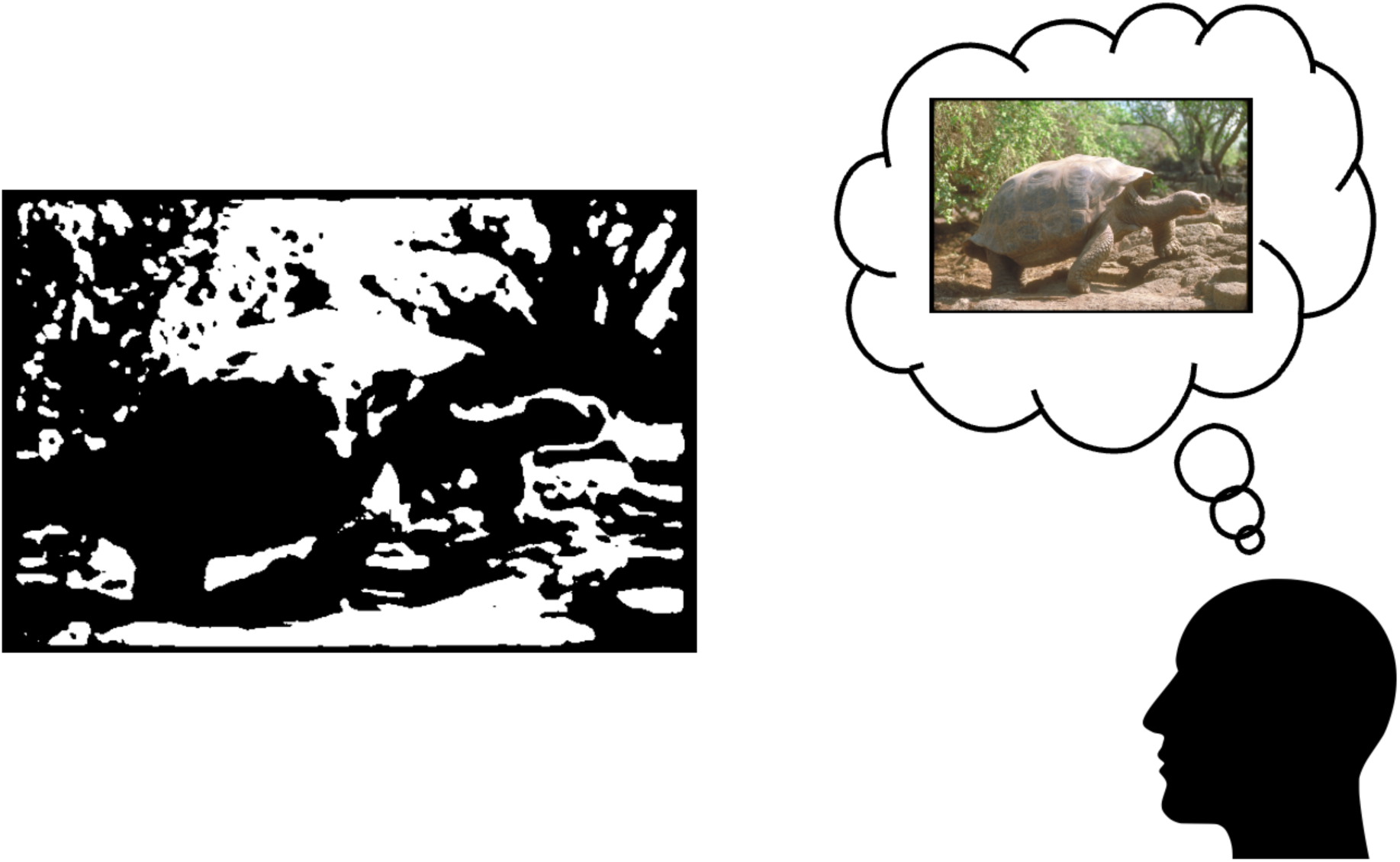
An Example of a Mooney Figure. *Note.* Observe the Mooney image on the left. At first glance, recognising its hidden content is challenging. Now, direct your attention to the original image inside the thought bubble. You may notice a dramatic shift in your perception upon revisiting the Mooney image. This phenomenon demonstrates how conceptual knowledge — acquired from the original image — guides the interpretation of latent visual structures. Our study leverages this unique property of Mooney images to examine whether object representations maintained in working memory can enhance perception by refining low-level visual details.

The simplicity and ambiguity of Mooney images make them ideal for testing how higher-level object knowledge influences low-level perception because they require the observer to fill in missing visual information, putatively via a top-down process (Mooney, 1957). Indeed, Mooney images demand top-down processing because their abstract, high-contrast format lacks clearly defined object features, making it close to impossible to recognise without prior learning (González-García et al., 2018; González-García & He, 2021; Mooney, 1957). Given these unique properties, and inspired by Teufel et al. (2018), the present study leverages Mooney images to investigate whether maintaining object memories enhances the refinement of low-level visual perception.

## The Current Study

We aimed to investigate whether holding a complex object concept in memory influences low-level perception. Specifically, we tested if observers’ detection sensitivity is influenced by object memory maintenance. In our experiment, each observer was shown a Mooney image that they held in working memory. During the memory retention interval, an edge-feature target appeared in 50% of trials. This differs from Teufel et al.’s approach, where participants were asked to detect the presence of targets within the Mooney images themselves. By shifting the focus to detecting features after the disappearance of the Mooney images, we aimed to isolate the effects of object memory maintenance on subsequent visual processing. To test if changes in visual sensitivity depend on the inferred contents of memory, the edge feature was either aligned to an implied border in the remembered Mooney image or rotated 90° (see Figure 2). By having observers perform this task before and after learning the hidden content of Mooney images, we manipulated whether observers held in memory the basic visual features versus conceptual information, respectively (Clarke & Tyler, 2015; Linde-Domingo et al., 2019). Additionally, we implemented attentional checks throughout the experiment to ensure that participants did indeed hold the Mooney image in working memory. We hypothesised that participants would have greater perceptual sensitivity to edge-defined targets embedded within Mooney stimuli after learning to discern the concealed objects. This heightened sensitivity was anticipated to be more pronounced for targets aligned with the implied edges of the learned objects, as opposed to those orthogonal.

**Figure 2.**
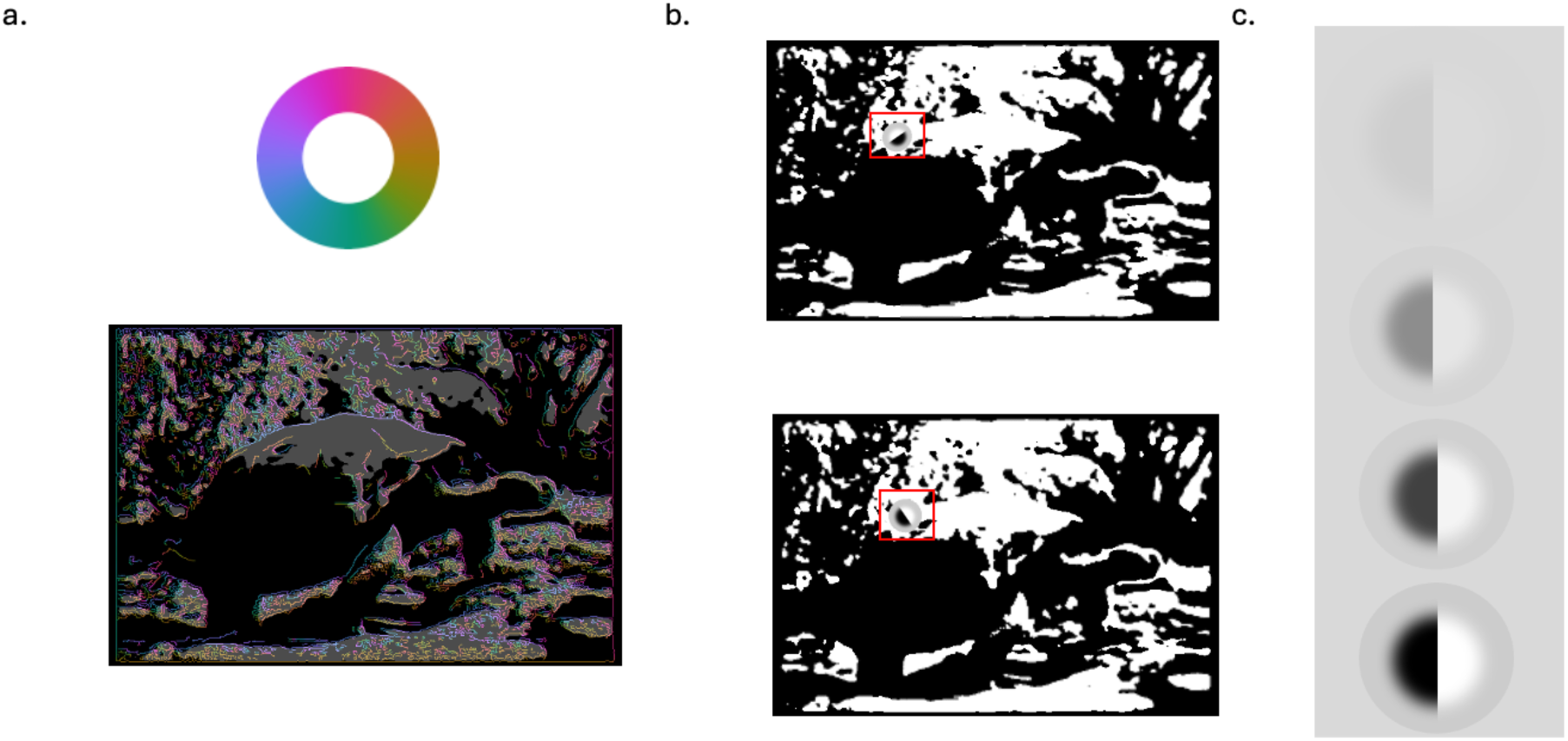
Examples of Alignment and Contrast Manipulation. *Note.* Example of Mooney image stimuli and target manipulations. (a) We used a steered filter approach to find oriented features along the border of hidden content (see Methods). (b) Examples of the same target, but either aligned with or orthogonal to the hidden image content. Aligned target (top): A Mooney image with a superimposed edge-feature target (highlighted in red) aligned with the underlying object contours. Orthogonal target (bottom): The same Mooney image with an edge-feature target rotated 90 degrees clockwise, disrupting alignment with the object contours. Note that, in the experiment, the targets were presented on a blank background, after the offset of the Mooney image. (c) Representation of the varying contrast levels used for target detection, shown along a low to-high contrast continuum. These examples are illustrative; the experimental contrasts were selected according to each observer’s contrast threshold.

## Method

### Participants

Twelve participants were recruited through the University of Queensland’s paid research portal. Two were excluded, failing to generate reliable psychometric functions within a maximum of two attempts (see below). Of the remaining ten (aged 21 to 25), nine were female and reported normal or corrected-to-normal vision. They spent an average of one and a half hours in the experiment and received $10 (AUD) compensation for every half-hour. The current study was approved by the university’s ethics committee.

### Materials

We used the Mooney stimuli from the Teufel et al. (2018) study, which consisted of forty-eight images centred around object categories such as still tools, animals, and people in natural scenes. We chose this dataset because these images were assessed based on a rigorous evaluation process with high convergent validity involving multiple judges (see Teufel et al., additional materials for further details). This stimulus set included the original photos, as well as the Mooney counterparts. Both types of images were used in different phases of the experiment. The original image was used to reveal the latent image content in its Mooney partner when teaching observers about the images (see below), as well as for computing all possible target orientations. The Mooney images were used during the experiment. For all possible target positions within a Mooney image, we used the steered filter approach outlined by Rideaux et al. (2022) to extract the orientations from the original images. In brief, this step involved convolving 0° and 90° oriented derivative-of-Gaussian filters with each image, and then combining the outputs of the filters to estimate local orientation.

For each image, experimenters (the first and senior authors) agreed on a single target location based on the visual information most relevant for recognition of the Mooney image. This was necessary because the image processing method described above is naïve to the contents of the image, computing orientations at all edge locations regardless of whether the edge is relevant to the primary object in the image. Therefore, target locations were first manually selected along the edges of implied contours (i.e. where there was oriented contrast in the original image but zero contrast in the corresponding Mooney image). Having selected the target location in each image, a perceptual target could then be steered to the local orientation at the chosen location or made to appear 90° orthogonal to local orientation. The target was an edge feature made from two opposite halves of 2D isotropic Gaussians with opposite polarities (see Figure 2c). As a result, it appears as a sharply defined edge even though it shares the same mean luminance as the background. In other words, the image creates an impression of a distinct boundary without any overall brightness difference between the two adjoining regions. This design creates the impression of a well-defined line while allowing us to manipulate local contrast changes as intended.

### Procedure

The current study used a blocked trial procedure, consisting of five sessions conducted in a fixed sequence. Participants provided written consent at the beginning of the experiment and were then instructed to watch a brief video explaining the Mooney images, as well as the keyboard and mouse controls for the experiment. Pre-testing sessions were conducted to select five Mooney images (session 1) and four contrast levels (session 2) tailored to each participant, based on their self-reported clarity of perception of Mooney images (before training) and baseline sensitivity in edge-feature target detection. The first two sessions establish the psychophysical baseline against which each individual will be evaluated in the main experiment

In the main experiment, participants viewed displays of Mooney images and performed a target detection task. Between the "before learning" session (session 3) and the "after learning" session (session 5), a learning session was interleaved (session 4), which included a knowledge test to ensure participants had learned the material successfully. Following this, participants completed another round of clarity ratings for the Mooney images (session 5). Participants completed all sessions in the same order, resulting in a repeated-measures experiment with a 5 (Mooney image) × 4 (target contrast) × 2 (session: before learning vs after learning) design.

#### Pre-testing Sessions

##### Thresholding Contrast Levels

Before the main experiment, we calibrated the contrast of the targets for each individual to ensure that we selected contrast levels around each participant’s threshold. During this calibration, participants viewed target stimuli flashed inside a fixation box for 200 milliseconds and used the left or right arrow keys to indicate whether they detected a target (left for “I saw a target”, right for “I did not see a target”). We used the QUEST method within Psychtoolbox-3 to measure participants’ target detection accuracy across various contrast levels (Watson & Pelli, 1983). Unlike the traditional method of constant stimuli in which a fixed set of predetermined contrast levels is presented repeatedly, the QUEST method adjusts the stimulus intensity based on participants’ responses. This adaptive procedure allows for an efficient determination of each participant’s contrast sensitivity threshold by setting the point where detection performance reaches a specific criterion. For the current experiment, we used a 75% correct response as the criterion.

##### Selection of Contrast Levels

Our a priori stimulus design aimed to include targets with four contrast levels, spanning two standardised deviations above and below each participant’s threshold in logarithmic space. This approach ensures that each observer views targets ranging from below-threshold contrast to above-threshold contrast. However, in cases where a participant’s contrast threshold was exceptionally low (i.e., below a minimum threshold of 0.005), an alternative approach was necessary to prevent non-negative values and adhere to the luminance limits of our 8-bit monitor. For participants with a measured threshold lower than 0.005, we assigned the first contrast level as 0.005. The subsequent contrast levels were selected based on linear increments, enabling us to assess sensitivity to relatively low contrast while ensuring that the task remained challenging yet feasible.

##### Selection of Mooney Stimuli

To select five target Mooney images for each individual, participants viewed all forty-eight Mooney images and rated their ability to perceive the “hidden” objects on a scale from 1 to 7, with a higher number indicating a clearer perception. The five images that received the *lowest* clarity ratings were chosen for further testing. Notably, while the study by Teufel et al. (2018) selected twenty out of forty-eight images, we chose a more conservative approach by focusing on only the five most ambiguous images reported by each observer. This adjustment was intended to emphasise the most challenging stimuli, providing a sharper contrast between initial perceptual difficulty and any potential improvement after the learning session. At the end of all sessions, participants repeated the rating procedure to determine whether their understanding of the hidden objects had improved. This post-experiment assessment focused on the five initially selected images with the lowest clarity ratings. By having observers repeat this Mooney rating at the end of the experiment, we could confirm whether they learned the hidden content by testing if their ratings increased relative to before the experiment.

#### Main Experimental Sessions

At the start of each trial, a randomly selected Mooney image flashed briefly for 500 milliseconds, followed by a red box indicating the target location (Figure 3). The target was then displayed for 200 milliseconds at the highlighted location — or omitted in 50% of the trials. Participants were instructed to press the left arrow key if they did not see a target (absent) or the right arrow key if they did (present). This main experimental session consisted of 300 trials and was conducted twice, once before, and once after participants learned about the hidden objects (see *Mooney Learning and Attention Checks* below).

**Figure 3.**
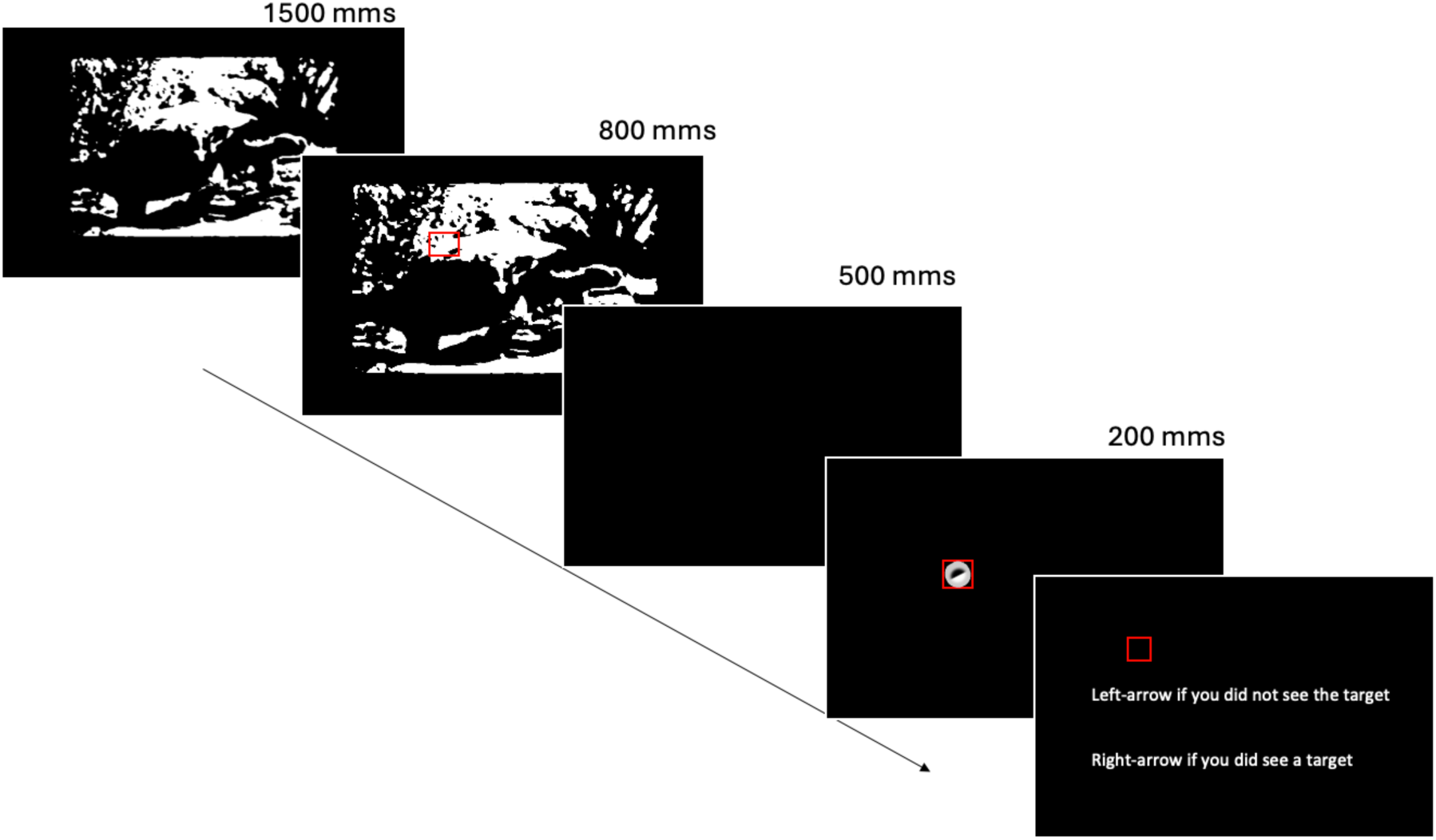
Trial Sequence for the Main Experiment. *Note.* Experimental trial sequence for target detection in Mooney images. (Left to right): Participants were first shown a Mooney image for 1500 milliseconds. A red box then indicated the target location, and a target stimulus was displayed inside the box for 200 milliseconds. Participants then indicated whether they saw the target using the left arrow key if no target was detected or the right arrow key if a target was present.

#### Mooney Learning

We revealed the latent structure in Mooney images only after a participant completed the first block of testing. To teach participants about the latent structure, we employed a *two-step* learning procedure like that of Teufel et al. (2018). In the first step, participants viewed the pre-selected Mooney images as they gradually faded in and out of their original forms over three cycles. This progressive revelation was designed to help participants develop an initial understanding of the hidden objects. In the second step, participants were prompted to closely examine the visual details of the original images. Specifically, they were instructed to track a moving red dot using a mouse cursor. During a single trial, the dot slowly ramped up from mid-level contrast, moving from the centre to the corners of each image in a random direction and distance. Successful tracking concluded with a mouse click at the final location where the red dot landed. Participants repeated this tracking task five times for each of their pre-selected images.

Following these steps, we tested if a participant had learnt the latent structure using a knowledge test. In this test, participants were shown a Mooney image and had to select from a list of four semantic labels, which label was most appropriate for the Mooney image. As an example, a participant may have been shown a Mooney version of a turtle image shown in Figure 1 and was required to select an appropriate label from one of four labels, including “tiger”, “whale”, “cat”, and “turtle” (shown in random order). Any incorrect response triggered a repeat of the fade-in-fade-out demonstration for that image. We repeated this semantic test halfway through the “after learning” main experimental session. This repetition reinforced participants’ semantic understanding and helped verify their retention of the meaningful content inside the images.

#### Attention Checks

To ensure participants were holding the Mooney images in memory, we implemented attention checks during the main experiment sessions. These checks, administered randomly across trials, aimed to prevent participants from focusing solely on the target detection task while ignoring the Mooney images. During each attention check, participants completed a forced-choice task at the end of a trial. They were asked to identify the Mooney image presented in the previous trial from two options: the correct image and a distractor image. The distractor image was randomly chosen from the remaining four Mooney images assigned to each participant. To ensure participants actively remembered the hidden objects, these checks were introduced in both the before and after learning main experimental sessions without prior notice. However, when they appeared on the screen, clear text instructions guided participants through the task: “Which of the two images did you see in the previous trial? Use left and arrow keys to respond”.

Notably, in the "after learning" phase of the main experimental session, incorrect responses during an attention check triggered a temporary pause in the experiment. During this pause, the corresponding original image was displayed again in a fade-in-fade-out sequence. This reinforcement was designed to strengthen participants’ memory of the hidden objects and help them maintain focus throughout the experiment.

### Main Experiment Analysis

A generalised linear mixed model (GLMM) was conducted to formally investigate the effects of object memory on low-level perception. The model was fit to participants’ responses (coded as 0 for absent, 1 for present) under the manipulated conditions, including sessions (before and after learning), four contrast levels, and the alignment of edge-feature targets. We did not include the five Mooney images as a factor in the current model (AIC: 5368), as the addition of this as a random effect did not result in a better model fit (AIC: 5370). Because the task design was a single interval yes/no task (Macmillan & Creelman, 1991), the GLMM included as a predictor the target’s absence or presence. Using a probit link function, the weights associated with the interactions between the target code and the different experimental factors are equivalent to *d’* (i.e. sensitivity) across conditions.

We were primarily interested in understanding whether object memory influences perception. Therefore, we expected the testing session (before vs. after learning the latent Mooney image content) to interact with each of the other manipulated factors. Interactive terms were incorporated to assess the influence of contrast, target alignment, and session. Participants’ IDs were treated as random effects to account for individual differences in detection performance.

The full model is expressed as follows:

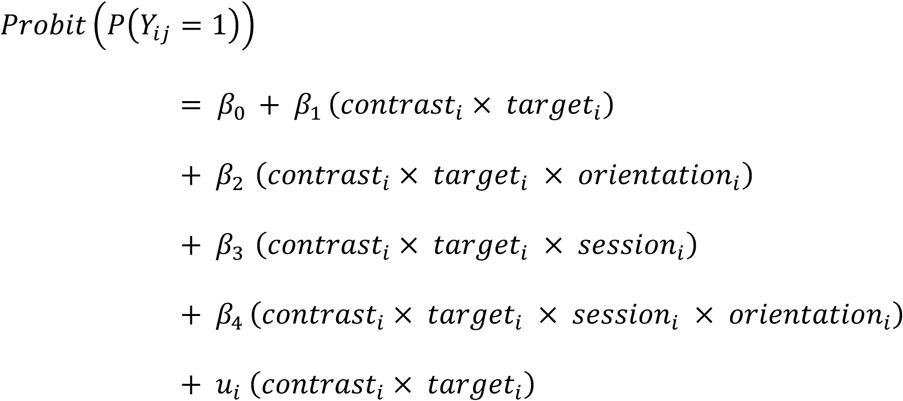

Here, the dependent variable is modelled as the probability of reporting the presence of the target. The *β* coefficients weight:

*β*_0_: The intercept, representing the baseline probability of reporting target presence regardless of the true state of the target. This is directly analogous to the signal detection theoretic response bias.

*β*_1_: The effect of the interaction between contrast and target presence, capturing how response probability depends on whether the target is present or absent and the contrast of the target.

*β*_2_: The effect of the three-way interaction among the contrast, target presence, and orientation alignment, because we predict that response probability also depends on whether targets are aligned or not with the latent Mooney structure.

*β*_3’_: The effect of the three-way interaction among contrast, target presence, and session, which assesses whether detection performance changes after learning the latent Mooney structure.

*β*_4_: The effect of the four-way interaction among contrast, target presence, orientation alignment, and session, capturing how these factors collectively influence detection probability.

Finally, the symbol *u_i_* denotes that we included a random slope for the interaction between contrast and target presence in the GLMM, accounting for individual differences in sensitivity to contrast levels across participants.

## Results

### Pre-testing Sessions

We measured individuals’ performance in detecting edge-feature targets to select an appropriate range of contrast levels for the main experiment. In this task, observers were required to respond whether they saw an edge-feature target or not. Plotted in Figure 4a are each observer’s psychometric functions. Participants’ ability to detect the targets did not vary substantially (mean threshold = 0.026, SEM = 0.005). In addition to finding each observer’s threshold, we also screened Mooney images, so the set of five images used for each participant depended on which images initially appeared most ambiguous. The five Mooney images with the poorest rating were highly variable across participants, as depicted in Figure 4b.

**Figure 4.**
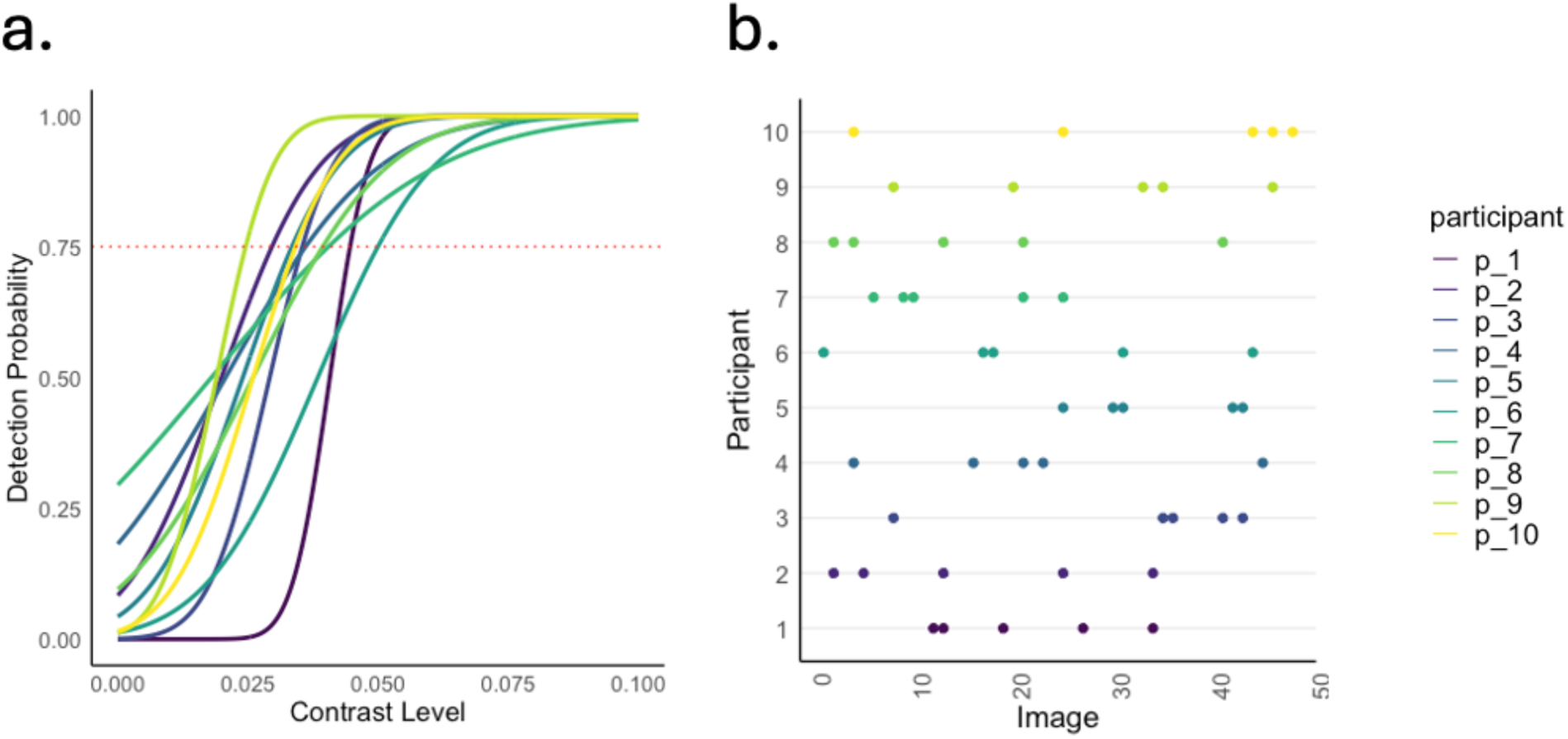
Individual Differences in Contrast Sensitivity and Mooney Image Screening. *Note.* (a) Detection probability across contrast levels for ten participants (p_1 to p_10). Each curve represents the psychometric function for individual participants, with the red dashed line indicating a detection threshold at 0.75 probability. (b) Individual responses across the full set of forty-eight images for the same ten participants show the variability in how they perceive Mooney images. Colours represent different participants across both plots.

### Semantic and Attention Checks

Our primary aim was to investigate how perception is influenced by memory maintenance of inferred visual structure. To test this hypothesis, we first need to establish 1) that participants learned the latent Mooney image structure, and 2) that they maintained this structure in memory during the perceptual task.

To test whether participants learned the latent image structure, we had them rate the subjective clarity of the Mooney images before and after learning the latent content. Should they learn the latent image structure, their subjective clarity ratings would be greater after learning than before. Consistent with this expectation, participants’ ratings of image clarity were higher after learning than before learning (*t (49)* = −14.40, *p* < 0.001; Figure 5). We therefore conclude that participants learned and perceived the latent image structure.

**Figure 5.**
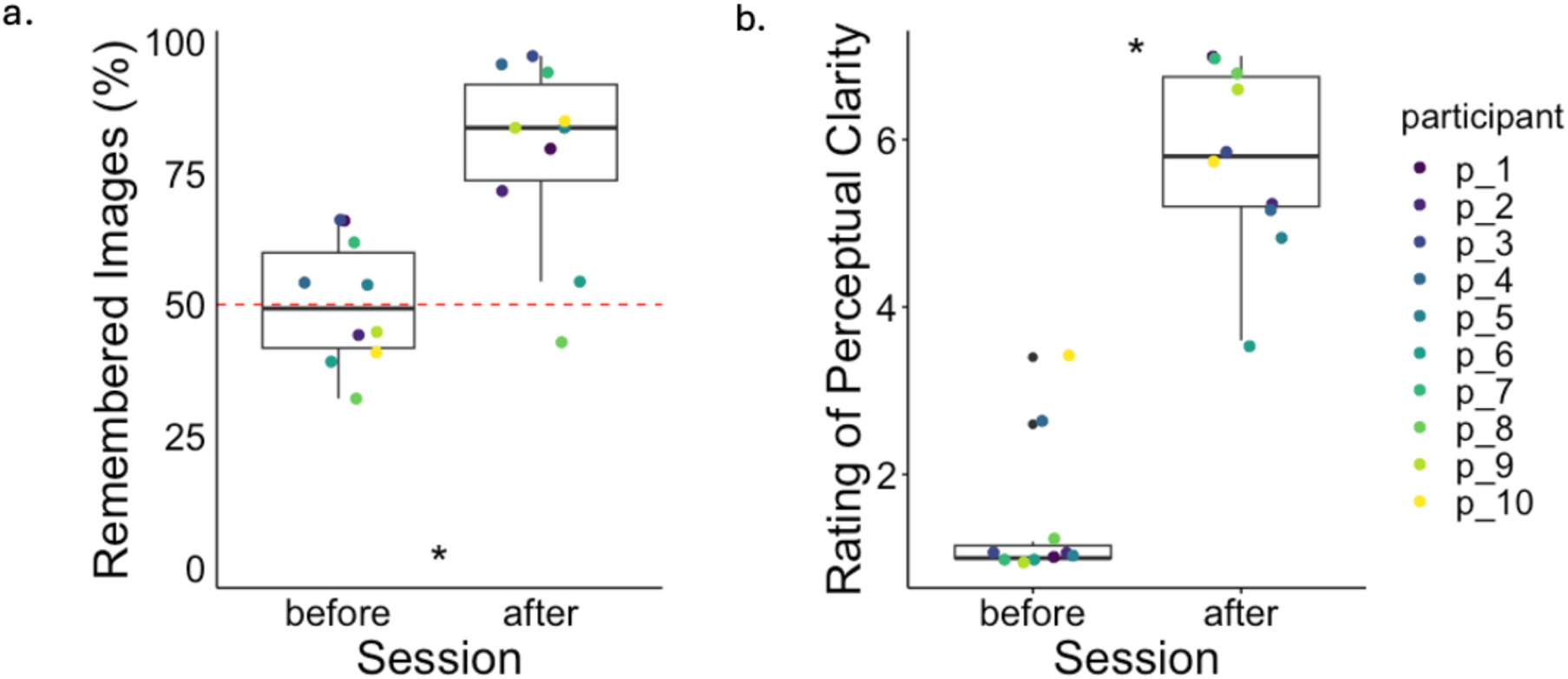
Results of Semantic and Attention Checks. *Note.* The percentage of correct judgement in attention tasks as a function of the session (before vs. after) showed a significant increase in the ability to remember which one of the two Mooney images was presented in the previous trial after learning. The red dashed line indicates the baseline for 50% accuracy; our data showed that participants barely performed greater than this baseline in session one before learning. (b) Average rating of subjective perception as a function of session (before vs. after), illustrating an overall increase in the perceived clarity of Mooney images. Significant differences between sessions are indicated by an asterisk (*).

To motivate participants to maintain the memory of the Mooney image throughout the perceptual test, we tested their memory of the previously shown Mooney image at the end of a subset of randomly chosen trials (see *Attention Checks* in Methods). If participants’ accuracy is at chance, we cannot be sure that they were holding the Mooney image in memory while making their responses. Before learning the latent content of Mooney images, participants’ ability to recognise the previously shown Mooney image was no different to chance (M = 50.28%, *t* = 0.076, *p* = 0.941). By contrast, their mean accuracy was 78.73% after learning the latent image content, which was a significant improvement (*t* (9) = −7.63, *p* < 0.001; see Figure 7). Therefore, participants were no better than chance before learning, but performed well after learning. This result demonstrates that participants were indeed holding the Mooney image in memory during the perceptual task of the after-learning testing session.

These results provide good evidence that participants learned the latent structure in the Mooney images, and, after learning, held the image content in memory during the perceptual task. We next test whether this memory maintenance of inferred structure influences perceptual sensitivity.

### Target detection sensitivity

We used a generalised linear model to quantify changes in sensitivity based on experimental manipulations, including session, target alignment, and contrast level. The model predictions, derived from a continuous range of contrast levels, represented the probability of responding "yes" to the presence of a target (regardless of whether the target was truly present or absent). These probabilities were then used to compute d’, capturing participants’ ability to discriminate between target-present and target-absent trials. This approach ensures that the sensitivity metric accounts for the discrimination of both the signal (target present) and noise (target absent) distributions independently of any individual’s response bias.

#### Target Sensitivity Before Learning

Before learning the hidden objects in the Mooney images, participants’ sensitivity to edge-feature targets depended strongly on contrast level, *β* = 0.59, SE = 0.12, *p* < 0.001 (see Figure 6). There was no difference in sensitivity to targets that were aligned versus misaligned with the contours of the hidden objects, *β* = −0.01, SE = 0.03, *p* = 0.62. This lack of difference was expected: participants had not yet learned about the hidden objects within their selected Mooney images, meaning their perception of targets ought not to be influenced by the memory of content embedded in the Mooney images.

**Figure 6.**
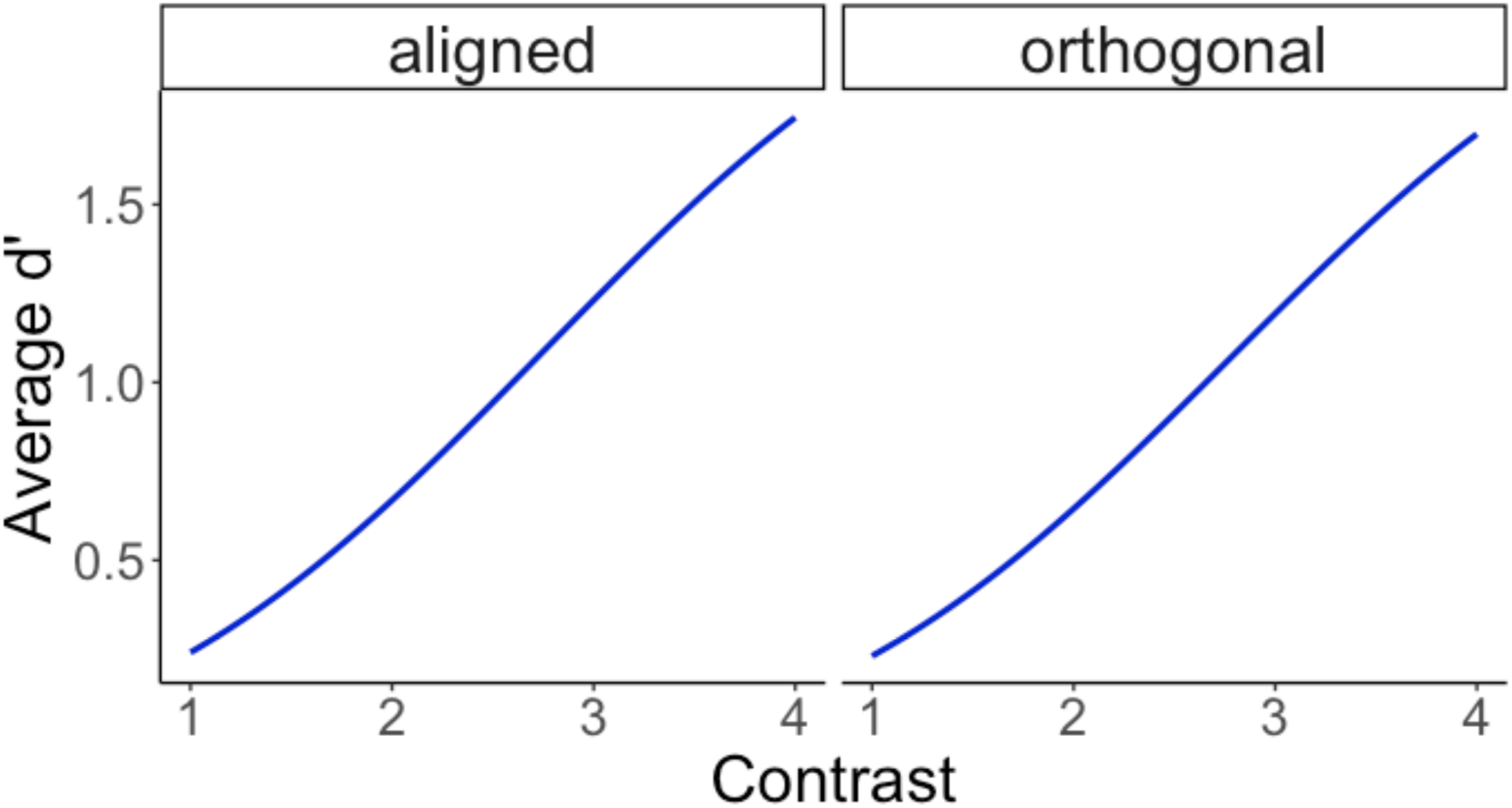
Sensitivity Towards Edge-feature Targets Before Learning the Mooney Image Structure. *Note.* Average predicted d’ across contrast levels for two different orientations (aligned in red and orthogonal in cyan). The plot shows a similar increase in d’ for both orientations with increasing contrast, indicating that the orientation of stimuli has minimal effect on the contrast sensitivity before learning.

#### Target Sensitivity After Learning

The critical test of whether learning the hidden content changed target sensitivity was assessed by the interaction between target presence, contrast, and testing session. This interaction was not significant, *β* = 0.02, *SE = 0.03, p = 0.40*. It could be that this analysis failed to find an influence of memory on perception because it averages over aligned and orthogonal target types. However, including the alignment of the target in the model did not change the result: the four-way interaction between contrast, target, orientation, and session was non-significant, *β* = −0.01, SE = 0.04, *p* = 0.83. Together, these findings suggest that, contrary to our initial hypothesis, learning about the latent structure in Mooney images did not significantly influence low-level perception (Figure 7). There are multiple possible reasons for this lack of an effect, which we discuss below.

**Figure 7.**
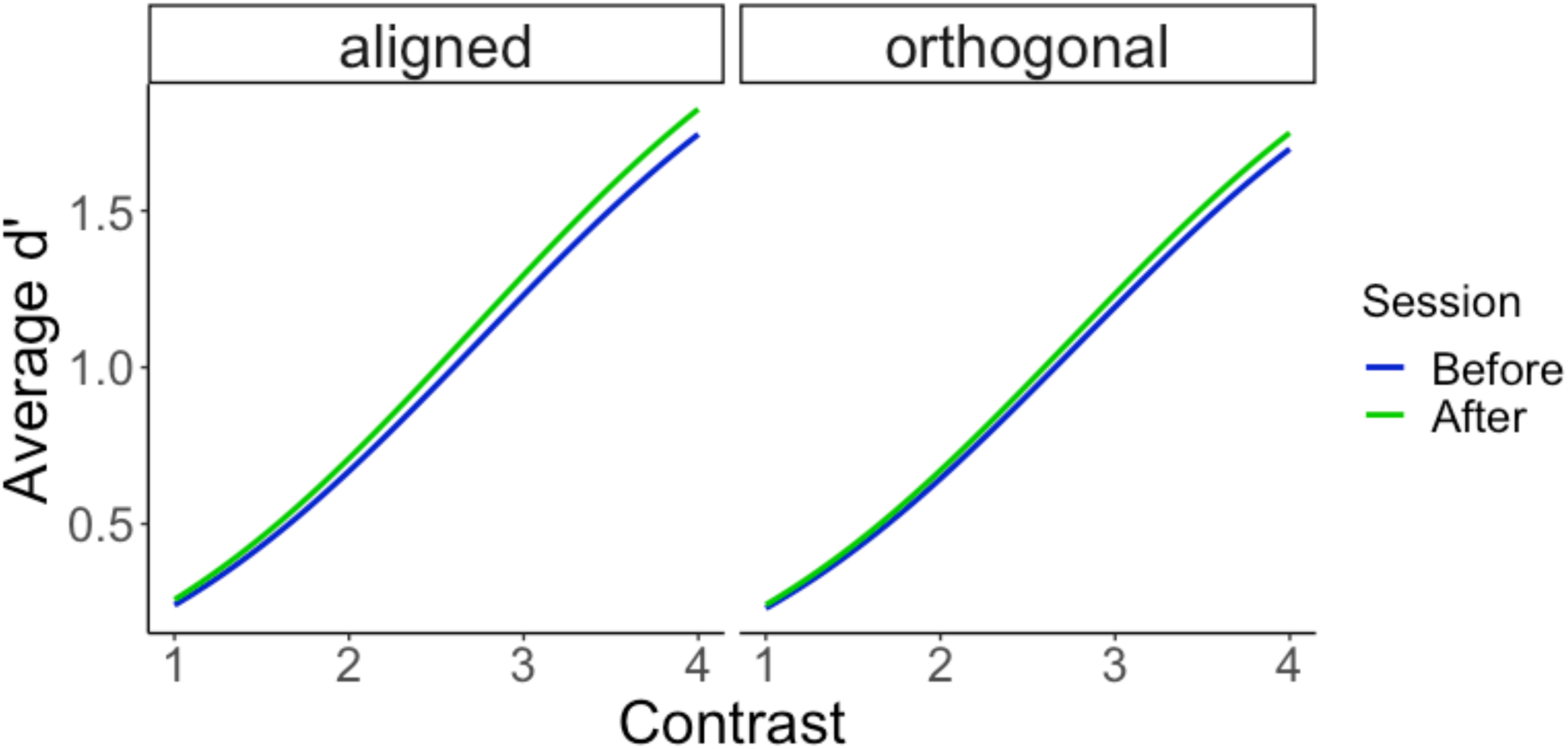
A Comparison of Before and After Learning Sessions. *Note.* Average predicted d’ as across contrast levels for two sessions (Session before in red and Session after in cyan) across two stimulus orientations: aligned and orthogonal. The plots show a consistent increase in d’ for both orientations, with minimal differences between sessions, suggesting stable performance across sessions for each orientation.

## Discussion

Recurrent processing is the notion that perception is shaped via the continuous refinement of low-level processing (Bar, 2004; Kveraga et al., 2007). It remains unclear, however, if visual representations maintained in working memory are sufficient to influence ongoing perception via a similar process to recurrent processing. Here, we combined the rigour of a psychophysics paradigm, incorporating signal detection theory, with the novel application of Mooney images to investigate whether inferred object memories influence ongoing perception by refining low-level visual details. Our participants successfully learned the latent content of Mooney images, resulting in substantial changes in their perception of the images (Figure 5). However, participants’ memory of inferred visual structure was not accompanied by a change in detection sensitivity of visual details associated with object memories. This suggests that, contrary to our prior expectations, object memory might not exert influence on perception through perceptual refinement. In the following discussion, we consider what this null effect implies for the popular theory, as well as possible reasons we may not have detected an effect that may indeed exist.

### Changes in Perceptual Judgement, not in Sensitivity, Reflect Categorical Processing

Using a signal detection framework, we disentangled the influence of perceptual sensitivity under varying conditions of decisional noise. Our results indicate that improved recognition did not coincide with enhanced sensory refinement. This key finding, while contrasting with proponents of recurrent processing, aligns with research suggesting that memory influences decisional confidence rather than altering sensory processing (Benwell et al., 2017; Bouyer et al., 2023; Rungratsameetaweemana et al., 2018; Summerfield & de Lange, 2014; Weaver et al., 2019).

For instance, Weaver et al. (2019) found that self-reported confidence ratings strongly predicted the disambiguation of binocular rivalry images (e.g., face vs. house), yet these ratings were unrelated to sensory representations in the early visual cortex. This suggests that complex object memory plays a crucial role in perception at a decisional level rather than influencing low-level sensory processes. Our results build on this perspective. While participants reported that the clarity of Mooney images increased after learning the latent image content, their sensitivity in detecting fine visual details did not improve. This dissociation suggests that complex object memory maintenance does not directly enhance low-level sensory processing. Interestingly, our participants verbally reported "seeing" the edge feature targets more clearly after learning, reinforcing the idea that their improved recognition of Mooney images reflects increased decisional confidence rather than sensory refinement.

Our study also extends the original findings of Teufel et al. (2018), whose paradigm inspired our research but produced different results. Teufel et al. reported that participants became more sensitive to edge-feature targets after learning Mooney images, suggesting that memory enhances perception by refining sensory details. In their paradigm, however, edge-feature targets were presented within the context of the Mooney image. In contrast, our paradigm assessed participants’ sensitivity to edge-feature targets immediately *after* viewing Mooney images. Taken together, it is possible that perceptual refinement occurs when viewing ambiguous images, but not when those images are held in memory.

Our findings diverge from neural studies that report activity in early visual regions during memory-guided perception (González-García et al., 2018; González-García & He, 2021; Johnson & Rugg, 2007; Kok et al., 2016; Kragel et al., 2021; Slotnick et al., 2005). The absence of psychophysical evidence in our study suggests that these findings may not reflect the refinement of sensory details proposed by the recurrent processing theory (Bar et al., 2006; Kveraga et al., 2007). While neural methods offer valuable insights, they suffer from correlational limitations. For instance, an enhancement in early visual activity may reflect memory maintenance (e.g., spatial locations of objects) (Ester et al., 2009; Hallenbeck et al., 2021; Summerfield & de Lange, 2014), attentional engagement (Boehler et al., 2008), or other higher-order processes rather than direct refinement in sensory representations (O’Connell et al., 2012; Summerfield & de Lange, 2014). By applying a signal detection framework with psychophysical testing, our study offers clear evidence of an instance in which memory-guided perception does not entail low-level sensory enhancement. This underscores the necessity of behavioural validation alongside neural measures to accurately interpret the mechanisms underlying perception and memory.

In summary, while memory-based expectations improve object recognition under uncertainty, our findings suggest they stem from categorical processing, not perceptual refinement. Complementing prior neural studies (Gwilliams & King, 2020; Linde-Domingo et al., 2019; Morales-Torres et al., 2024; Von Seth et al., 2023; Xie et al., 2024), our behavioural findings are consistent with a change in decisional confidence over sensory enhancement, but this remains to be confirmed in future experiments. Our findings highlight the importance of behavioural methods in a field dominated by neuroimaging, where distinguishing sensory from decisional influences remains a challenge (Summerfield & de Lange, 2014).

### Alternative Explanations and Current Limitations

Although our findings did not support our prediction that maintaining complex object memory influences low-level perception, we acknowledge potential caveats in our study design. These limitations may have prevented us from detecting the hypothesised effect, particularly because the specific paradigm and stimulus selection may be critical in uncovering such effects. One notable limitation is our relatively small sample size, which may have underpowered the critical statistical analyses. To address this concern, we conducted a Bayesian version of the GLMM, which provides the level of evidence for the hypothesised effect, taking into consideration the sample size (see Supplementary Material). This analysis produced similar results and further strengthened our conclusions. Specifically, the Bayesian model indicated some moderate evidence favouring the alternative hypothesis but also showed substantial uncertainty. Future research with larger samples will help validate and refine these findings. Nevertheless, if perceptual refinement were fundamental to recurrent feedback, a small sample size alone would not have obscured its detection. The fact that we found no evidence for perceptual refinement suggests that the effect may be negligible, even if it does exist. The following sections discuss alternative explanations for these results and offer recommendations for future experiments.

#### Individual Differences Might Modulate Perceptual Refinement

Reverse correlation methods, as employed by Canas-Bajo and Whitney (2022), have revealed idiosyncratic perceptual templates that appear to reflect "fill-in" sensory refinement for each individual participant in perceiving Mooney faces. This is consistent with recognition memory literature, which emphasises that object concepts can be stored in different representational formats depending on individual differences and motivational factors (Wilson et al., 2012). Since our study focused on pre-selected edge feature details (one feature per image); it may not fully capture the diverse perceptual cues participants might have used to interpret Mooney images. The absence of individual feature selection in our study may have contributed to the null result, as participants might have relied on perceptual cues beyond the pre-selected edge features.

Future research could improve upon our study by allowing participants to identify which edge features are most salient to them, potentially revealing more nuanced individual differences in perceptual refinement if it exists. Further research using adaptive paradigms that account for individual differences in feature selection could clarify this issue. For example, a recently developed psychophysical technique that manipulates uncertainty in Mooney image generation offers a promising avenue for investigating how individual perceptual strategies shape the interpretation of ambiguous stimuli (Reining & Wallis, 2024). Integrating such approaches could provide deeper insights into how memory-driven perception varies across individuals.

#### High-level, Not Low-level, Perceptual Integration

Alternatively, our observed lack of effect may align with recent evidence suggesting that perceptual refinement primarily involves higher-order features, while low-level features remain unchanged but continue to support the recognition process (Gwilliams & King, 2020; Mohsenzadeh et al., 2018; Morales-Torres et al., 2024; Von Seth et al., 2023; Xie et al., 2024; Zhao et al., 2024). Within this framework, representations in the early visual cortex act as a cognitive interface during recurrent feedback, interacting with higher-order processes like memory to guide more complex representations flexibly (Bracci et al., 2017; Rademaker et al., 2019; Xie et al., 2024; see Roelfsema & De Lange, 2016 for review).

For example, Rademaker et al. (2019) showed that early visual regions maintain stable sensory representations even under interference, providing a robust foundation for comparing incoming sensory inputs with memory-based representations. Recent studies using time-sensitive methods further revealed that, although the early visual cortex re-engages during recurrent feedback, semantic information is decoded for mid to high-level features (Morales-Torres et al., 2024; Von Seth et al., 2023; Xie et al., 2024). In a similar vein, research by Gayet et al. (2016, 2018) showed that memory enhances perception by lowering the threshold for evidence sampling rather than increasing sensory gain. Thus, instead of modifying existing low-level representations, recurrent feedback likely enables flexible interpretation by emphasising task-relevant, high-level information (Bracci et al., 2017; Rademaker et al., 2019; Zhao et al., 2024). This distinction is critical for maintaining a balance between constructing a functional reality and avoiding the overreliance on inferences that could lead to hallucinations (Calderone et al., 2013).

Although we did not directly measure changes in mid or high-level features, our psychophysical findings contribute to prior research by demonstrating improvements in categorical processing without alterations in low-level sensory details. Future studies may therefore benefit from targeting higher-order features inside Mooney images to investigate whether such features indeed drive perceptual refinement. Mid-level visual features in Mooney images, like the turtle shell, might be extracted early on but only fully integrated once participants tap into their knowledge of what a turtle looks like (Clarke & Tyler, 2015; Vandenbroucke et al., 2014). Examining how different levels of perceptual cues interact with memory-driven interpretation could clarify whether prior knowledge enhances the ability to extract meaningful structure from ambiguous stimuli. For instance, real-world object forms and sizes can be reliably detected from shape or texture alone, even after controlling for low-level cues (Long et al., 2016).

Finally, integrating psychophysical approaches with time-sensitive neural methods (Gwilliams & King, 2020; Mohsenzadeh et al., 2018; Von Seth et al., 2023) could provide deeper insights into how high-level perceptual integration unfolds over time. While neuroimaging studies have highlighted early visual cortex involvement, a critical next step is determining how this activity contributes to categorical perception rather than low-level sensory refinement. By bridging behavioural and neural evidence, future studies could refine our understanding of how the visual system leverages memory to enhance recognition without altering fundamental sensory representations.

## Conclusion

Inspired by recurrent processing theory, we investigated whether object memories enhance conscious perception by refining low-level visual details using a novel Mooney paradigm. While object memories improved the perception of hidden objects, they did not enhance low-level visual information. Instead, our findings suggest that memory shapes perception by guiding overall understanding rather than refining sensory information. This highlights a flexible role for memory in perception, prioritising efficient integration of existing knowledge to support meaningful interpretations of sensory inputs.

## Supporting information

Supplementary Analysis

## Notes

### Competing Interest Statement

The authors have declared no competing interest.

