## Supplementary Analysis for "Do Memories of Inferred Visual Representations Guide Low-level Perception?"

### Bayesian Results

We conducted a Bayesian equivalent of the GLMM described in the study. Several key considerations informed this decision. First, contextual effects in similar studies are typically small, making it essential to utilise prior distributions to improve parameter estimation and guide the analysis. Second, the Bayesian framework is more closely aligned with the theoretical principles of predictive coding, which is important for our hypothesised top-down influence of memory on perception. For example, learning the contents of Mooney images is expected to adjust participants' prior expectations, thereby increasing the posterior probability of target detection. Finally, given our relatively small sample size (N = 10), the Bayesian model provides a level of evidence on a continuous scale, which differs from the GLMM presented in the main text that dichotomises the results as significant or not.

#### Model Parameterisation

Our model assumes that the response variable follows a Bernoulli distribution, where the probability of a positive response is $P\ (response = 1)$ modelled as:

$$P(response = 1 \mid predictors) = \ \Phi\ (\eta)$$

The $\Phi\ (\eta)$ indicates the cumulative distribution function (CDF) of the standard normal distribution, the probit link function. The bracketed $\eta$ reflect the linear predictors of the response variable, which are defined as:

$$\eta = \ \beta_0 + \ \beta_1 (contrast_i \times \ target_i) + \beta_2 \ (contrast_i \times \ target_i \ \times orientation_i) + \ \beta_3 \ (contrast_i \times \ target_i \ \times \ session_i) \ + \ \beta_4 (contrast_i \times \ target_i \ \times session_i \ \times \ orientation_i) + \ u_i \ + \ \epsilon$$

The response variable $P\ (response = 1)$ is modelled as a Bernoulli random variable with the predicted probability given by the probit function. Like the GLMM described in the Method section, the beta coefficients $\beta$ here also capture the interaction effects between contrast, target, orientation, and session. The term $u_i$ represents the random effect for each participant, allowing the model to account for individual variability and expressed in a normal distribution:

$$u_i \sim \mathcal{N}\,(0,\ \sigma^2)$$

**Prior Implementation**

For the four-way interaction, we specified normal priors with a mean of 0.8 and a standard deviation of 0.5, reflecting a moderate positive effect based on the results of Teufel et al. (2018). Contrast-related coefficients were assigned priors with a mean of 3 and a standard deviation of 2, capturing our strong expectation for targets with higher contrasts to be detected compared to those with lesser contrast. As for the rest of the interaction coefficients, we applied general priors with a mean of 0.5 and a standard deviation of 0.5, introducing a mild positive interaction while maintaining narrower uncertainty.

**Model Results**

The Bayesian model estimated the response probability using contrast, target presence, orientation, and session as predictors, interacting with these factors. The model converged successfully ($\hat{\mathcal{R}}$ < 1.01) without divergent transitions. Posterior predictive checks demonstrated that the model fit the data well, accurately capturing the observed patterns.

As shown in Table 1 below, the contrast has a significant and positive main effect on the probability of a response, $\beta$ =1.46, 95% CI [ 0.57, 2.35]. This result suggests that increased contrast levels resulted in an increased probability of participants reporting the presence of the targets, as expected. As for interaction terms, their effects are minimal and less certain, with many credible intervals including zero. This indicates that higher-order interactions may have limited or negligible contributions under the conditions tested (see Table below).

A closer examination of the Bayes factors revealed that there might be moderate evidence for an interaction between contrast and session, BF = 4.66. However, for the higher-order interaction involving contrast, target presence, orientation and session, there was little support for the hypothesised effect, BF = 0.68. Taken together, these results highlight the central role of contrast in influencing detection responses. While some interaction terms show moderate evidence for their effects, the evidence for contributions of higher-order interactions is equivocal.

**Table 1**

*Regression Coefficients of the Bayesian Model*

| Regression Coefficients | Estimate | Est.Error | 95% CI | 95% CI | Rhat |
|---|---|---|---|---|---|
| Intercept | -0.7 | 0.29 | -1.26 | -0.1 | 1 |
| contrast:target | 1.46 | 0.46 | 0.57 | 2.35 | 1 |
| contrast:target:orientation | -0.03 | 0.12 | -0.26 | 0.2 | 1 |
| contrast:target:session | 0.11 | 0.12 | -0.12 | 0.34 | 1 |
| contrast:target:orientation:session | -0.04 | 0.16 | -0.36 | 0.28 | 1 |

**Model Predictions**

Model predictions were derived from posterior estimates to compute d' (d-prime) metrics — a formal measure of the complex hierarchical relationships within the dataset. As illustrated in Figure 1a, participants' sensitivity to targets increased slightly after learning, with aligned targets demonstrating a more pronounced improvement. This pattern aligns with the findings of Teufel et al. (2018), supporting the reliability of our results. Figure 1b shows no changes in false alarm rates across either orientation condition following learning. Finally, Figure 1c on hit rates revealed a pattern of increase like d' across both orientations between sessions. Together, these findings provide nuanced insights into how sensitivity and response patterns might shift because of learning, emphasising the distinct effects for aligned targets.

Recall that the regression coefficients of the Bayesian model did not strongly support these seemingly substantial interaction effects. The transformation of the d-prime calculation process may introduce additional nonlinearity, resulting in visually compelling results. On the one hand, d prime metrics provide a straightforward analysis for understanding changes in decision-making and perceptual sensitivity. On the other hand, the apparent interaction effects must be interpreted with caution due to the lack of robust statistical support in the original model.

**Figure 1**

*Model Predictions of Sensitivity (d prime), False Alarm and Hit Rates*

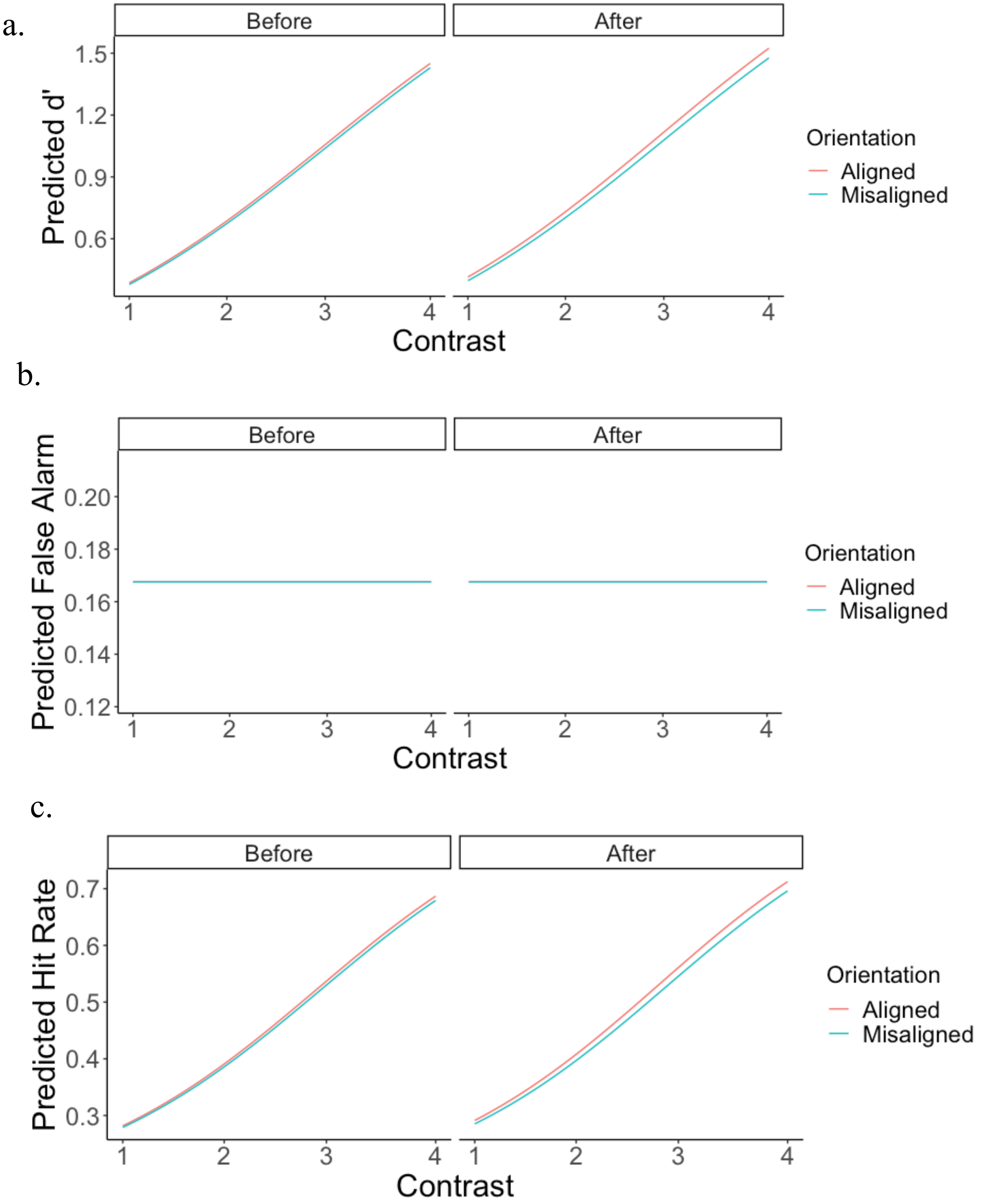
